# Dynamic bi-directional phosphorylation events associated with the reciprocal regulation of synapses during homeostatic up- and down-scaling

**DOI:** 10.1101/2021.03.26.437166

**Authors:** Kristina Desch, Julian D. Langer, Erin M. Schuman

**Author notes:** lead contact: erin schuman.

## Abstract

Homeostatic synaptic scaling allows for bi-directional adjustment of the strength of synaptic connections in response to changes in their input. Protein phosphorylation modulates many neuronal and synaptic processes, but it has not been studied on a global, proteome-wide scale during synaptic scaling. To examine this, we used LC-MS/MS analyses to measure changes in the phosphoproteome in response to up- or down-scaling in cultured cortical neurons over minutes to 24 hours. Out of 45,000 phosphorylation events measured, ~3,300 (associated with 1,280 phospho-proteins) were regulated by homeostatic scaling. The activity-sensitive phosphoproteins were predominantly located at synapses and involved in cytoskeletal reorganization. We identified many early transient phosphorylation events which could serve as sensors for the activity offset as well as late and/or persistent phosphoregulation that could represent effector mechanisms driving the homeostatic response. Much of the persistent phosphorylation was reciprocally regulated by up- or down-scaling, suggesting that the mechanisms underlying these two poles of synaptic regulation make use of a common signaling axis.

## Introduction

Adjustments to neuronal network properties can be achieved by activity-dependent plasticity. Homeostatic synaptic scaling is one such form of plasticity that involves the compensatory and global up- or down-regulation of synaptic strength in response to changes in the level of synaptic input. For example, homeostatic up-scaling occurs when network activity is reduced by the cessation of action potentials. In contrast, homeostatic down-scaling can be elicited when network activity is elevated by blocking inhibitory synaptic transmission (Turrigiano, 2012, 2008). Mechanistically speaking, both up-scaling and down-scaling converge on the modulation of synaptic AMPA receptors to bring about the compensatory change in synaptic strength (O’Brien et al., 1998; Turrigiano et al., 1998). Other molecules like NMDA-type glutamate receptors, postsynaptic scaffold proteins such as PSD-95 and Homer, or soluble factors like Bdnf have been implicated in synaptic scaling as have global changes in the synthesis and degradation of proteins (Dörrbaum et al., 2020; Ehlers, 2003; Schanzenbächer et al., 2016).

For scaling to occur, neurons must possess molecular mechanisms that sense the change in overall activity level (sensors) and then implement the scaling response (effectors). Both modeling and experimental studies suggest that intracellular Ca^2+^ levels may serve as an activity-regulated early signal during scaling (Ibata et al., 2008; Marder and Prinz, 2003; Thiagarajan et al., 2005). An important and ubiquitous post-translational modification, often Ca^2+^-sensitive, that can alter a protein’s catalytic activity, localization, interactions or stability is phosphorylation (Humphrey et al., 2015). Ca^2+^-sensitive kinases and phosphatases enable differential phosphorylation and dephosphorylation of various target proteins. For example, Ca^2+^/calmodulin-dependent protein kinase II α (Camk2a) is a well-studied kinase that is important for many different forms of plasticity. During the long-term potentiation (LTP) Camk2a undergoes increases in auto-phosphorylation on Thr^286^, resulting in Ca^2+^-independent kinase activity (Fukunaga et al., 1993; Miller and Kennedy, 1986). While phosphorylation events are perhaps ideal for the initial detection of plasticity-induced stimuli, the long-term regulation of kinases (including Camk2a) and phosphatases can also mediate persistent aspects of synaptic and behavioral plasticity (Farley and Schuman, 1991; Malinow et al., 1988; Martin et al., 1997).

Many synaptic receptors and scaffold elements are known to be phosphorylated in the context of normal synaptic function (Greengard et al., 1993; Nestler and Greengard, 1983) and plasticity (Diering et al., 2017; Engholm-Keller et al., 2019; van Gelder et al., 2020; Kohansal-Nodehi et al., 2016; Lee, 2006). Data from a previous broad-scale proteomics analysis (Schanzenbächer et al., 2016) revealed kinases and phosphatases as the largest differentially regulated group of newly synthesized proteins during homeostatic scaling (Schanzenbächer et al., 2016). Differential protein phosphorylation during synaptic scaling has also been described by candidate-based approaches (Jang et al., 2015; Sanderson et al., 2018). Recently, Yong *et al*. described the phosphorylation of the AMPA receptor subunit 2A (Tyr^876^) during the late phase of homeostatic up-scaling using phospho-deficient knock-in mice (Yong et al., 2020). Other recent studies have examined phosphorylation during sleep (Brüning et al., 2019; Diering et al., 2017), in short-term plasticity/stimulation (Engholm-Keller et al., 2019; van Gelder et al., 2020; Kohansal-Nodehi et al., 2016; Li et al., 2016) or studied its role in a disease-related context, e.g. Alzheimer disease (AD) (Bai et al., 2020).

Here, we investigated proteome-wide protein phosphorylation following global, bidirectional homeostatic scaling in primary cultured cortical neurons. We made use of a bottom-up, LC-MS-based proteomics pipeline that enabled us to identify and quantify unmodified and phosphorylated proteins in a global and unbiased manner. We monitored the activity-sensitive phosphorylation events from minutes to one day following scaling, establishing a dataset of ~3,380 differentially regulated phosphorylation events of ~1,280 unique proteins associated with different phases and types of homeostatic scaling. We found distinct regulatory signatures converging on proteins of the synaptic compartment which exhibited persistent, reciprocal and time-sensitive phosphorylation. A quarter of the initial phosphorylation events detected were persistently regulated in a reciprocal manner reflecting the opposite polarity of up- and down-scaling.

## Results

### Activity-dependent protein phosphorylation in cortical neurons

To investigate the dynamics of activity-dependent protein phosphorylation in cortical neurons, we conducted MS-based phosphoproteomics during homeostatic plasticity (Figure 1A). Homeostatic up- or down-scaling was induced by treatment with Tetrodotoxin (1 μM; TTX) or Bicuculline (20 μM; Bic), respectively. We explored the temporal dynamics of protein phosphorylation during plasticity by examining protein phosphorylation after 5 min, 15 min or 24 hrs of stimulation. Temporal snapshots of the proteome and the phosphoproteome were acquired by high-resolution LC-MS/MS analyses using a bottom-up, label-free proteomics approach (see Methods and Tables S1 and S2). Each of the six different experimental conditions was analysed in four independent biological replicates which were injected in three technical replicate LC-MS/MS runs.

**Figure 1.**
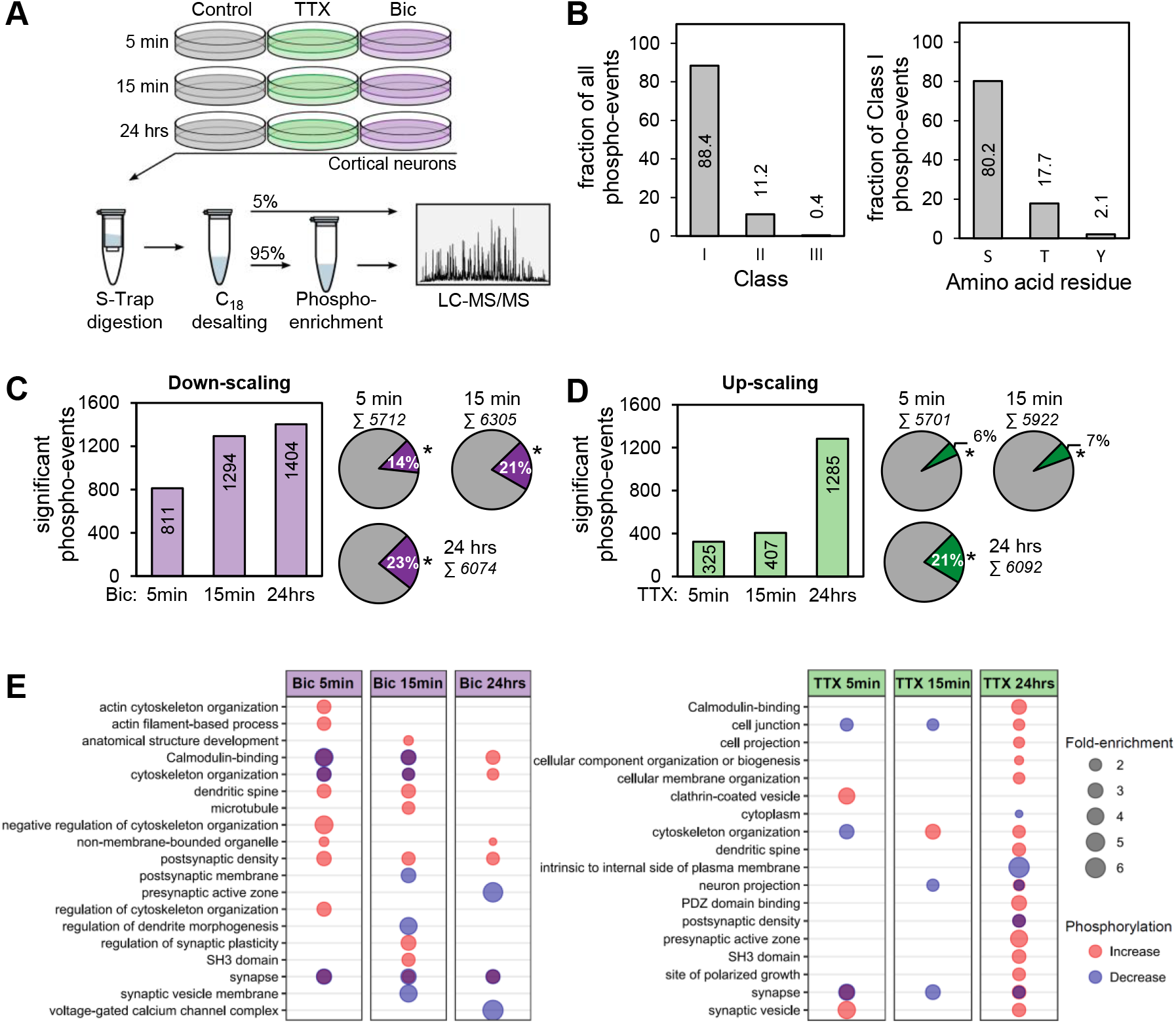
Quantitative LC-MS/MS analysis of activity-dependent phosphorylation in cortical neurons. (A) Illustration of the experimental workflow. Cultured cortical neurons (DIV 19-20) were treated with Tetrodotoxin (1μM; TTX) or Bicuculline (20μM; Bic) for 5 min, 15 min or 24 hrs. Subsequently, the cells were harvested and lysed. Digestion was performed using a suspension trapping protocol (S-Trap). After purification of the peptides, one part of the sample (5%) was directly investigated via LC-MS/MS analysis. A second part (95%) was retained for enrichment of phosphorylated peptides using TiO2-beads and then analyzed via LC-MS/MS (see Methods). For each of the six different conditions, four independent biological replicates were prepared which were measured in three separate LC-MS/MS runs. (B) Overview of the identified phosphopeptide species. The left graph shows the distribution of phosphorylated residues of all quantified phosphorylation events according to their localization probability (Class I: > 75% prob.; Class II: 50% to 75% prob.; and Class III:< 50% prob.). The right graph indicates the distribution of phosphorylated amino acid residues of all Class I phosphorylation events. (C and D). Temporal profile of phosphorylation events. The bar charts show the number of significantly regulated phosphorylation events (Benjamini-Hochberg correction; FDR < 0.01) comparing Bic (C) or TTX (D) treatment versus the control at each time point. The pie charts indicate the proportion of significantly (*) and not significantly regulated phosphorylation events. (E) Gene ontology (GO) overrepresentation analysis of phosphorylated proteins for all three time points of Bic (left) and TTX (right) treatment (Benjamini-Hochberg correction; FDR < 0.01). Proteins with a regulated phosphorylation site were divided with respect to the nature of the regulation (increase or decrease) prior to GO analysis. The degree of regulation is highlighted by the color and the fold enrichment is indicated by the size of the dots. If a term was enriched for both increased and decreased phosphorylation, the corresponding dot is shown in overlap and hence appears purple.

Across all experimental groups, our analysis detected 4,520 phosphorylated proteins identified in depth with 45,691 phosphopeptide species. These phosphorylation events were filtered for reliable phosphorylation site assignment with a location probability greater than 75% (Class I), allowing us to map the modification site to a specific amino acid residue (Figure 1B). In downstream analyses, peptides with insufficient modification site localization (Class II/III sites) were discarded. The Class I phosphorylation events had a depth of 40,395 peptide species, mapping to 26,642 unique phosphorylation sites. The distribution of phosphorylated serine, threonine and tyrosine residues was 80.2%, 17.7% and 2.1%, respectively (Figure 1B), similar to what has been observed in other proteomics experiments (Brüning et al., 2019; Ubersax and Ferrell, 2007). We quantified differential phosphorylation by comparing phosphorylated peptide species in each (up- or down-) scaling group to its associated control sample. Only phosphopeptide species passing stringent criteria were used in downstream analyses (see Methods).

The induction of homeostatic up- or down-scaling led to a large and global regulation of protein phosphorylation (see Table S3). Treatment with Bic (down-scaling induction) for 5 min, 15 min or 24 hrs resulted in a significant alteration of 811, 1,294 or 1,404 phosphorylation peptide species, respectively (referred to as “regulated phospho-events”; Figure 1C). Treatment with TTX (up-scaling induction) for 5 min, 15 min or 24 hrs resulted in a significant alteration of 325, 407 or 1,285 phosphorylation events, respectively (Figure 1D). The above changes in protein phosphorylation could result from plasticity-induced changes in protein levels, which have been previously documented (Dörrbaum et al., 2020; Schanzenbächer et al., 2016). We assessed the overlap of significantly regulated phospho-events with proteins that underwent significant changes in their abundance (Figure S1A-D, Table S4) and found a very small overlap (Figure S2A,B; 5%, 14% and 9% of the regulated phospho-events at 15 min Bic, 24 hrs Bic and 24 hrs TTX treatment were also regulated at the protein level).

The above dataset of 3,382 different activity-regulated phosphorylation events were associated with 1,285 unique proteins (Figure 1C,D; Figure S3). To reveal functional classes of the proteins significantly regulated by phosphorylation, we performed a gene ontology (GO) overrepresentation analysis of the regulated phosphoproteins (FDR < 0.01). The analysis was conducted separately for up- or downregulated phosphorylated proteins. Consistent with the change in synaptic strength elicited by scaling, we found overrepresentation of the term “synapse” during the whole course of stimulation for both up- and down-scaling indicating the differential phosphorylation of synaptic proteins (Figure 1E). For both up- and down-scaling there was also significant overrepresentation of “cytoskeleton organization” as the activity manipulation progressed. During Bic-induced downscaling, additional terms that were consistently overrepresented were “Calmodulin binding” and “postsynaptic density”. Altogether, the analysis revealed that during up- and down-scaling, there was rapid and long-lasting phosphoregulation in the synaptic compartment and a progressive re-organization of the cytoskeleton.

### Phosphoprotein dynamics during synaptic scaling

How is protein phosphorylation regulated over time during homeostatic scaling? For both up- and down-scaling, the extent of phosphoregulation clearly increased with the duration of the manipulation (Figure 1C,D). While most of the regulated phosphorylation events (62%, n = 2,112) were associated with a single time-point (down-scaling: 72% and up-scaling 84%; Figure S4), there were also a large number of persistent phosphorylation events (observed at all times) for each form of scaling (Figure 2A,B). To examine the regulation associated with the different phases of plasticity, we first binned the time-limited phospho-events for each type of scaling (Figure 2A, Table S5). A large number of transient events were detected in the early (n = 1,149) and late phase (n = 830) of down-scaling whereas during up-scaling the number of phosphorylation events increased dramatically over time (n= 392 and 1,061 for early and late time points). Most of the time-limited events were specific for either up- or down-scaling. These phosphorylation events could represent sensors/effectors that were specific to the sign of the plasticity. On the other hand, we also observed overlap in the phosphorylation targets observed in up- and down-scaling: 110 events during the early-phase and 149 during the late-phase of plasticity (Figure 2C). A large fraction of all overlapping regulated phosphorylation events was reciprocal in nature: positively regulated (increased) phosphorylation for one type of scaling and negatively regulated (decreased) in the other. These reciprocally regulated phosphorylation events could represent signaling pathways responsible for the detection of activity offsets from a set-point; the sign of the activity offset (increased or decreased) could then be represented by an increase or decrease in phosphorylation.

**Figure 2 –.**
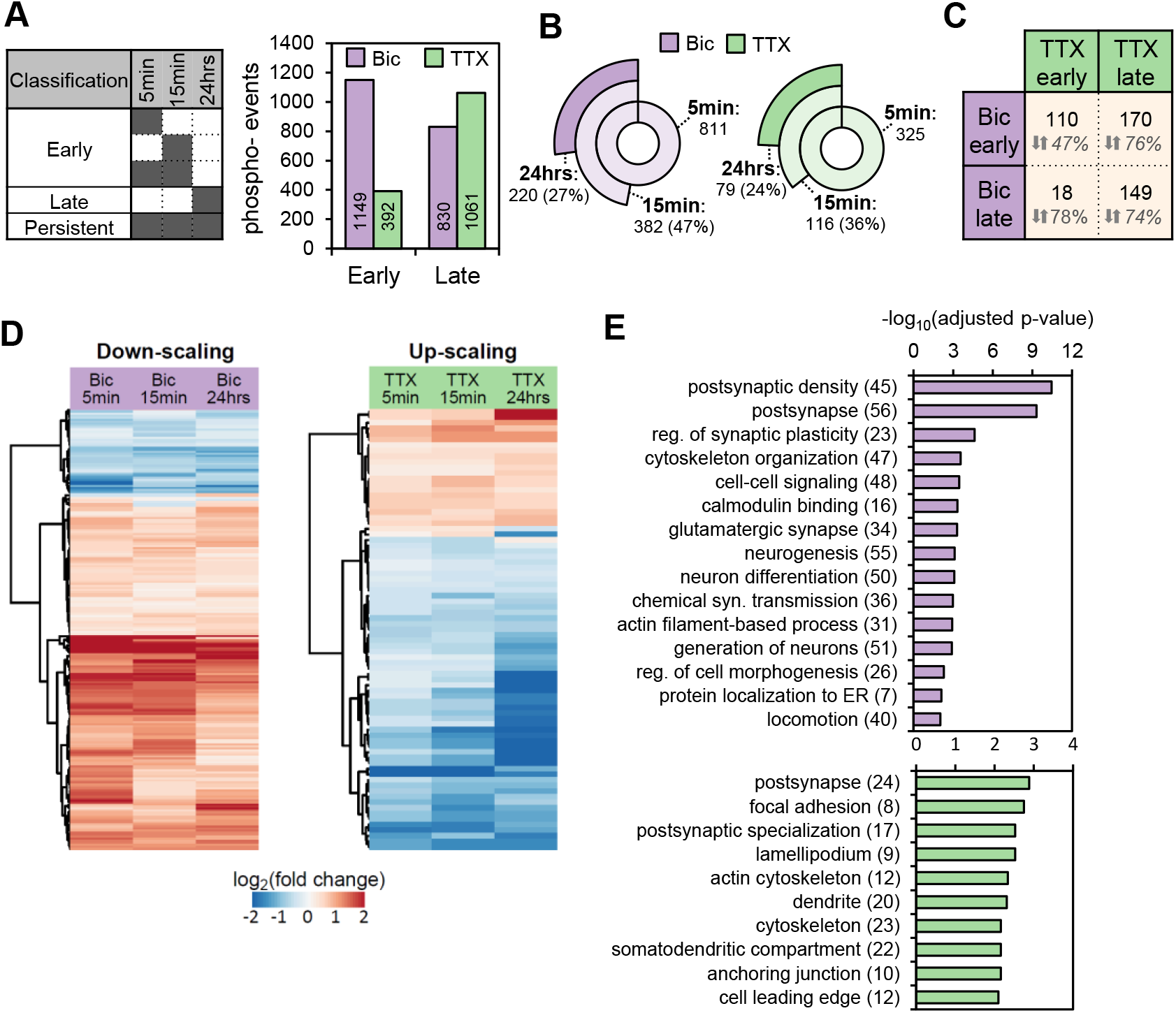
Phosphorylation dynamics during synaptic scaling. (A) Time-limited phosphorylation events were grouped according to the classification indicated in the table (left). Dark color indicates significant regulation at the highlighted time point. The bar chart (right) displays the distribution and number (insets on bars) of early and late phosphorylation events for Bic and TTX treatment. (B) Significantly regulated phosphorylation events over time. About a quarter (Bic: 27%; TTX: 24%) of the initially regulated phosphorylation events exhibited significant changes at all time points. (C) Overlap of the phosphorylation events in the temporal groups as observed between the treatment groups is indicated in absolute numbers in the table. The percentage of overlapping phosphorylation events exhibiting reciprocal regulation is written below. (D) Hierarchical clustering (Euclidean distance) was performed on the log_2_ fold changes (treatment vs. control) of the phosphorylation event intensities regulated during up- and down-scaling as described in (C). Shown is a heatmap (red: increase in phosphorylation, blue: decrease in phosphorylation) indicating two major clusters for both up- and down-scaling. There was a significant decrease in intensity over time in the down-scaling phosphorylation increase cluster and significant increase in intensity over time in the up-scaling phosphorylation decrease cluster (p < 0.05; ANOVA and Tukey HSD post hoc). (E) GO overrepresentation analysis of the persistent Bic-evoked (top) and TTX-evoked (bottom) subset of phosphorylation events (Benjamini-Hochberg correction; FDR < 0.01). Selected terms and their adjusted p-value is shown (see Table S6 for all terms).

To investigate the temporal evolution of phosphorylation during homeostatic scaling, we performed a hierarchical clustering analysis of the persistent phosphorylation events. Approximately a quarter of initial phosphorylation events (regulated at 5 min) exhibited significant regulation at all three tested time points (Figure 2B). We subjected the log_2_-transformed fold changes of all of the persistently regulated phosphorylation events to hierarchical clustering (Figure 2D; Figure S5). The long-lasting phosphorylation events formed two distinct clusters during up- or down-scaling; one cluster represented a persistent increase in phosphorylation and the other a persistent decrease in phosphorylation. During down-scaling, the phosphorylation increase cluster was large and the phosphorylation decrease cluster was smaller, whereas the opposite was true for up-scaling.

Which phosphorylated proteins are part of these persistently regulated clusters? We performed GO overrepresentation analyses for the persistently regulated phosphoproteins and found, again, many protein groups related to synaptic function (Figure 2E). For example, for both up- and down-scaling, the term “postsynapse” was overrepresented. Similarly, “postsynaptic density”, “regulation of synaptic plasticity” and “chemical synaptic transmission” were overrepresented in the Bic-induced persistent subset of signals and “postsynaptic specialization” in the TTX-induced persistent subset. The term “calmodulin binding” for phosphoproteins regulated during down-scaling was also observed. Indeed, we detected the persistent autophosphorylation site Thr^286^ on Ca^2+^/calmodulin-dependent protein kinase II alpha (Camk2a), which leads to Ca^2+^-independent enzyme activity (Miller and Kennedy, 1986). We also observed a persistent increase in the phosphorylation of Ser^421^ of Methyl-CpG-binding protein 2 (Mecp2), the Rett syndrome protein implicated in synaptic plasticity and an established Camk2a substrate (Amir et al., 1999; Zhou et al., 2006). Furthermore, phosphorylation at Ser^556^ on Synapsin 1 (Syn1), another Camk2a-regulated site (Czernik et al., 1987), also showed persistent regulation. Both lists of significantly overrepresented terms also included the cytoskeleton: “cytoskeleton organization” and “locomotion” during down-scaling or “actin cytoskeleton” and “cytoskeleton” during up-scaling.

### Comparing the phosphorylation pattern of up- and down-scaling

Homeostatic up- and down-scaling involve the reciprocal regulation of synaptic strength in response to activity that is either decreased or increased relative to a set-point. Although both forms of scaling converge on the same phenotypic endpoint (the number of synaptically localized AMPA receptors) the cell-signaling mechanisms are not well understood. To examine whether the same proteins were regulated by up- and down-scaling, we expanded the clustering analysis to phosphorylation events that were regulated in at least four of six possible conditions across both experimental groups. The phosphorylation events that were regulated in both scaling groups showed a remarkable reciprocity in the sign of regulation (Figure 3A): upregulated phosphorylation events during down-scaling were downregulated during up-scaling or vice versa. We generated protein-protein-interaction maps of the reciprocally regulated phosphoproteins of the two major clusters and annotated them (Figure 3B). The smaller cluster of reciprocal phosphorylation events was downregulated during down-scaling and upregulated during the up-scaling. Three small interaction-networks were formed by presynaptic proteins (Aak1, Bin1, Amph or Snap91), proteins associated with ubiquitin signaling (Nedd4l or Lmo7), but also by cytoskeletal proteins (Epb41l3 or Epb41l1). The second cluster of reciprocal phosphorylation events was upregulated during down- and downregulated during up-scaling. In total, nine interaction-networks were generated from this set. The smaller subnetworks comprise phosphoproteins directly involved in ubiquitin signaling (Uba1, Uba5, Ube2o or Herc1) or associated with translation initiation (Eif4b and Eif4g). Another subnetwork contained proteins of the presynaptic compartment following the same regulation pattern. The largest subnetwork comprised phosphoproteins of the postsynaptic compartment. Postsynaptic density protein 95 (Dlg4, PSD-95), a post-synaptic scaffold protein implicated in synaptic plasticity (Bats et al., 2007; Vallejo et al., 2017), emerged as a central hub. Other phosphoproteins that exhibited this reciprocal regulation profile were important cytoskeleton interactors such as Syngap1 or Map1a, but also proteins known to interact with or traffic neurotransmitter receptors (Grip1, Sh3kbp1 or Lrrc7). Of note, phosphorylation sites on some proteins in this network have been reported to affect the protein’s function directly, e.g. phosphorylated Ser^2798^/ Ser^2804^ of the Ryanodine receptor 2 (Ryr2) increases channel conductance (Ferrero et al., 2007; Huke and Bers, 2008).

**Figure 3 –.**
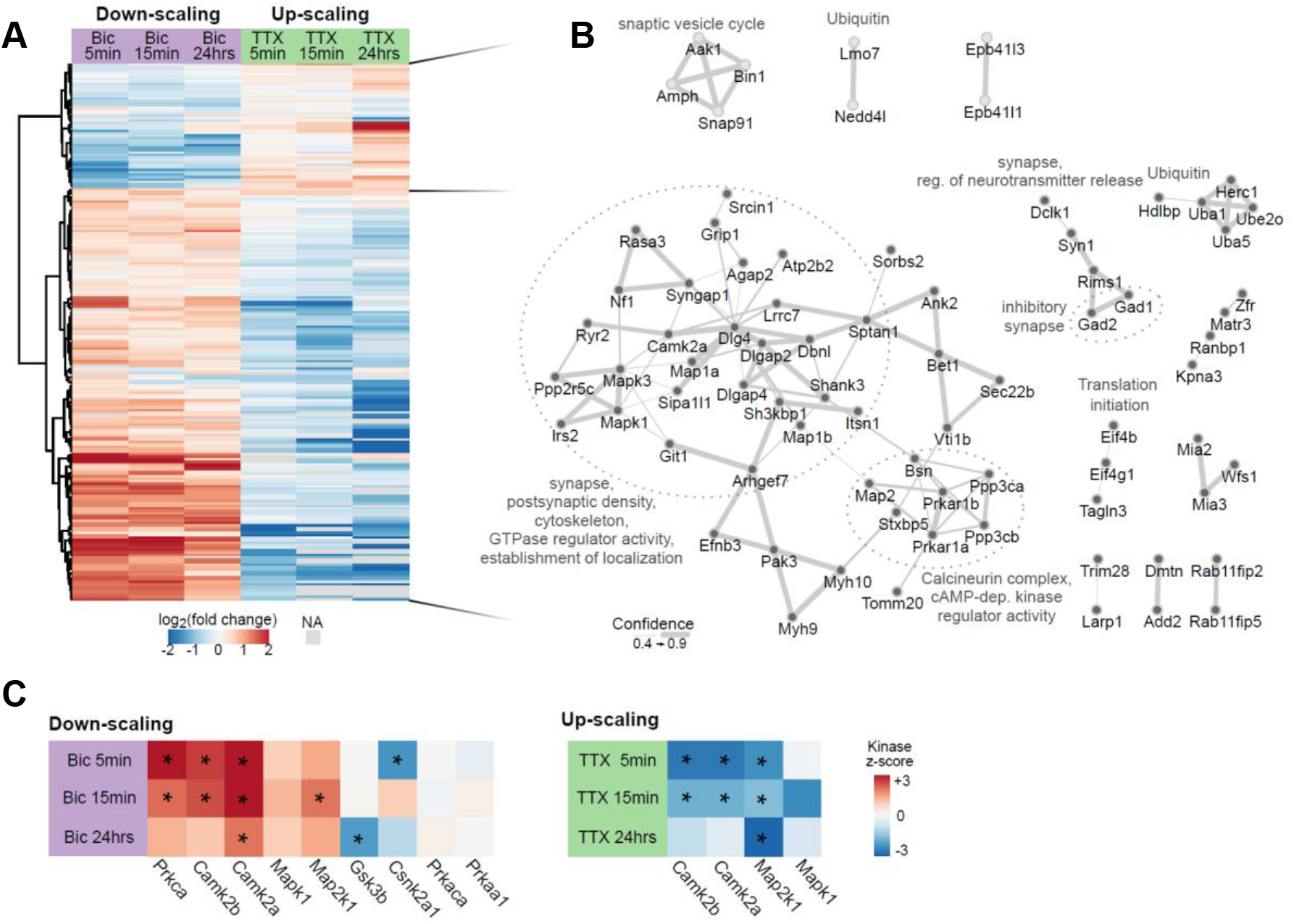
Persistent and reciprocally regulated phosphorylation patterns during up- and down-scaling. (A) Hierarchical clustering analysis shown with a heatmap combining both scaling experiments: the log_2_ fold change (treatment versus control) of phosphorylation events that were significantly regulated in at least 4 of 6 experimental conditions (n = 219) were clustered (Euclidean distance). Red: increase in phosphorylation, blue: decrease in phosphorylation. Missing values (NA): grey color. (B) Phosphoproteins of the two major clusters of the heatmap in (A) were analyzed for protein interactions using their gene identifiers and the String database. Cluster membership is indicated by the color of the nodes (light grey nodes = top cluster; grey nodes = bottom cluster). The sub-networks are described by overrepresented GO or manually curated terms (UniProtKB). Width of the connections show the confidence of the interaction (combined score) as derived from the String database. Unconnected phosphoproteins are not shown. (C) Prediction of kinase activity was performed via kinase substrate enrichment analysis (KSEA) on the significantly regulated phosphorylation events of each condition (see Methods). Kinase activity is visualized by color according to the kinase z-score (red: increase in kinase activity, blue: decrease in kinase activity). Significant regulation is highlighted by an asterisk within the tile of the heatmap (*; p <0.05, FDR <0.1).

To understand how this bi-directional phosphorylation patterns is achieved, we focused on the kinases that were differentially phosphorylated by scaling. Prominent activity-dependent kinases were central nodes in the postsynaptic subnetwork. For example, the activating phosphorylation sites of Camk2a (Thr^286^), ERK1 (Thr^203^/ Tyr^205^) or ERK2 (Thr^183^/ Tyr^185^) showed significant reciprocal regulation. We next performed kinase-substrate-enrichment analysis (KSEA) where kinase activity (kinase z-score; Figure 3C, Table S7) was calculated from the phosphorylation status of reported substrates (Figure S6). There, persistent and bi-directional behavior could be detected by the kinase z-scores of Camk2a and MAP-kinases.

We validated the patterns of Camk2a and MAP-kinase phosphoregulation using protein-specific antibodies together with phospho-specific antibodies (Figure 4). Overall, the pattern of phosphoregulation observed with phospho-specific antibodies was very similar to that observed in the MS data. Using immunoblotting, we found that down-scaling resulted in a significant increase in Camk2a Thr^286^ phosphorylation at both 5 min and 24 hrs whereas upscaling resulted in a trend for enhanced phosphorylation at 24 hrs (Figure 4A, B). For both ERK1 Thr^203^/ Tyr^205^ and ERK2 Thr^183^/ Tyr^185^ down-scaling resulted in a significant increase in phosphorylation at both 5 min and 24 hrs whereas upscaling resulted in a trend for enhanced phosphorylation at 24 hrs (Figure 4C, D). In contrast, upscaling resulted in a significant decrease in phosphorylation, which was a clear trend at 5 min and significantly different from control at 24 hrs (Figure 4C, D). Taken together, these data suggest that the activity of Ca^2+^-sensitive kinase Camk2a and ERK1/2 drive persistent phosphorylation during Bic-induced down-scaling while they appear deactivated during TTX-induced upscaling.

**Figure 4 –.**
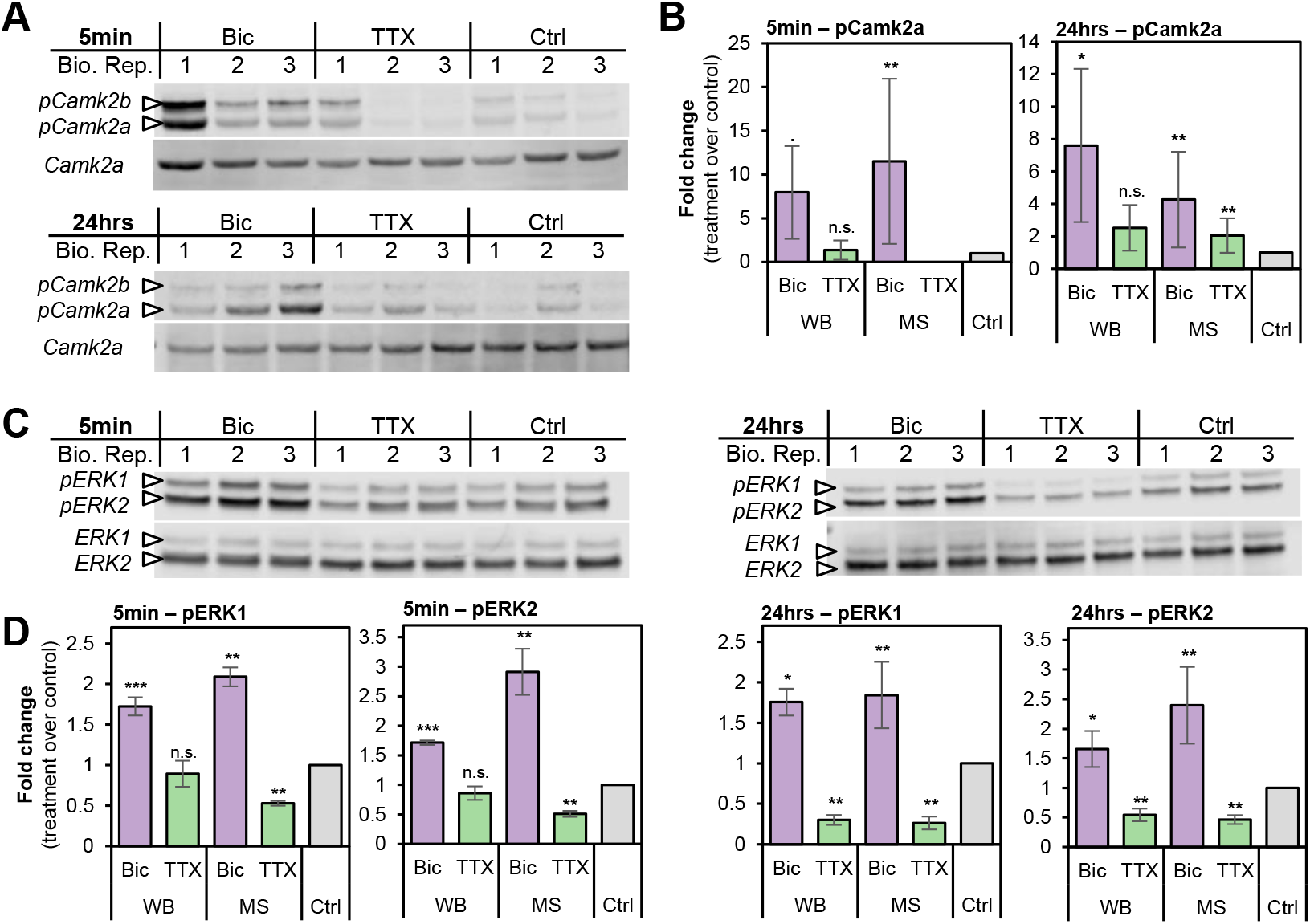
Verification of phosphoregulated Camk2a, ERK1 and ERK2 by immunoblotting. (A) Western blot images for pCamk2a (pThr^286^) and total Camk2a. Analyses were performed using three independent biological replicates of Bic-, TTX-treated and untreated (control) neurons for 5 min (top) or 24 hrs (bottom). (B) Bar graphs show quantification of pCamk2a detected by western blot (WB) in comparison the phosphopeptide’s signals quantified via LC-MS/MS (MS) after 5 min (left) or 24 hrs (right) Bic- or TTX-stimulation. Protein signals of the blot were quantified based on the intensity ratio of the phosphorylated protein over total protein normalizing treated conditions to the untreated control. Log_2_-scaled MS-intensity of the phosphorylated peptide is depicted as the fold change of treatment over control as well (see Methods for details). Error bars represent the standard deviation between biological replicates (Western Blot: n = 3; MS: n = 4). For statistical analysis of the western blot data, a two-sided t-test was performed: *** p>0.001; ** p<0.01; * p<0.05;. p<0.1; n.s., not significant. For MS-data, ** FDR <1%. (C) Western blot images for pERK1 (pThr286/pTyr205) or pERK2 (pThr183/pTyr185) and total ERK1 and ERK2. Analyses were performed using three independent biological replicates of Bic-, TTX-treated and untreated (control) neurons for 5 min (left) or 24 hrs (right). (D) Bar graphs show quantification of pERK1 and pERK2 detected by western blot (WB) in comparison the phosphopeptides quantified via LC-MS/MS (MS) after 5 min (left) or 24 hrs (right) Bic- or TTX-stimulation. Western blot and statistical analyses were performed and visualized as described in (B).

### Mapping the regulation of the synaptic phosphoproteome over time

To examine the different phospho-patterns in the synaptic compartment, we generated time-resolved and site-specific maps of the synaptic phosphoproteome during homeostatic up- and down-scaling (Figure 5, Table S8). The protein-based visualization of phosphoregulation affirmed the similarity between both scaling polarities, as seen by the common phosphoproteins (n = 94, marked with asterisks). Site-specific annotation revealed that different modes of phosphorylation converged on phosphoproteins in both the pre- and postsynaptic compartments. Many phosphoproteins were phosphorylated at more than one phospho-site in the course of the experiment. An example is the differential phosphorylation of Dlg4. During Bic-induced down-scaling, Dlg4 exhibited upregulation at three different sites (Tyr^240^, Ser^418^, Ser^422^) assigned to all three temporal classes (early, late, persistent/reciprocal). During TTX-induced up-scaling, significant changes were limited to three different phosphosites that were either continuously regulated (Ser^422:^ downregulated) or regulated during the late phase of stimulation (Ser^73^: upregulated, Ser^418^: downregulated). These modifications were mostly reciprocally regulated (Ser^422^, Ser^418^) and located within or close to the PDZ-domains of Dlg4; regions which have already been associated with fine-tuning Dlg4’s molecular associations (Pedersen et al., 2017). Another example of a multiple phosphorylation site protein is Rims1, a scaffold element at the presynaptic active zone that regulates neurotransmitter release (Castillo et al., 2002). Rims1 contained six regulated phospho-sites during down- and nine regulated sites during up-scaling. During the late phase of up-scaling, an increase in Ser^592^ phosphorylation of Rims1 (Ser^413^ in mice) has been observed, which is associated with LTP and the recruitment of 14-3-3 adaptor proteins (Simsek-Duran et al., 2004).

**Figure 5 –.**
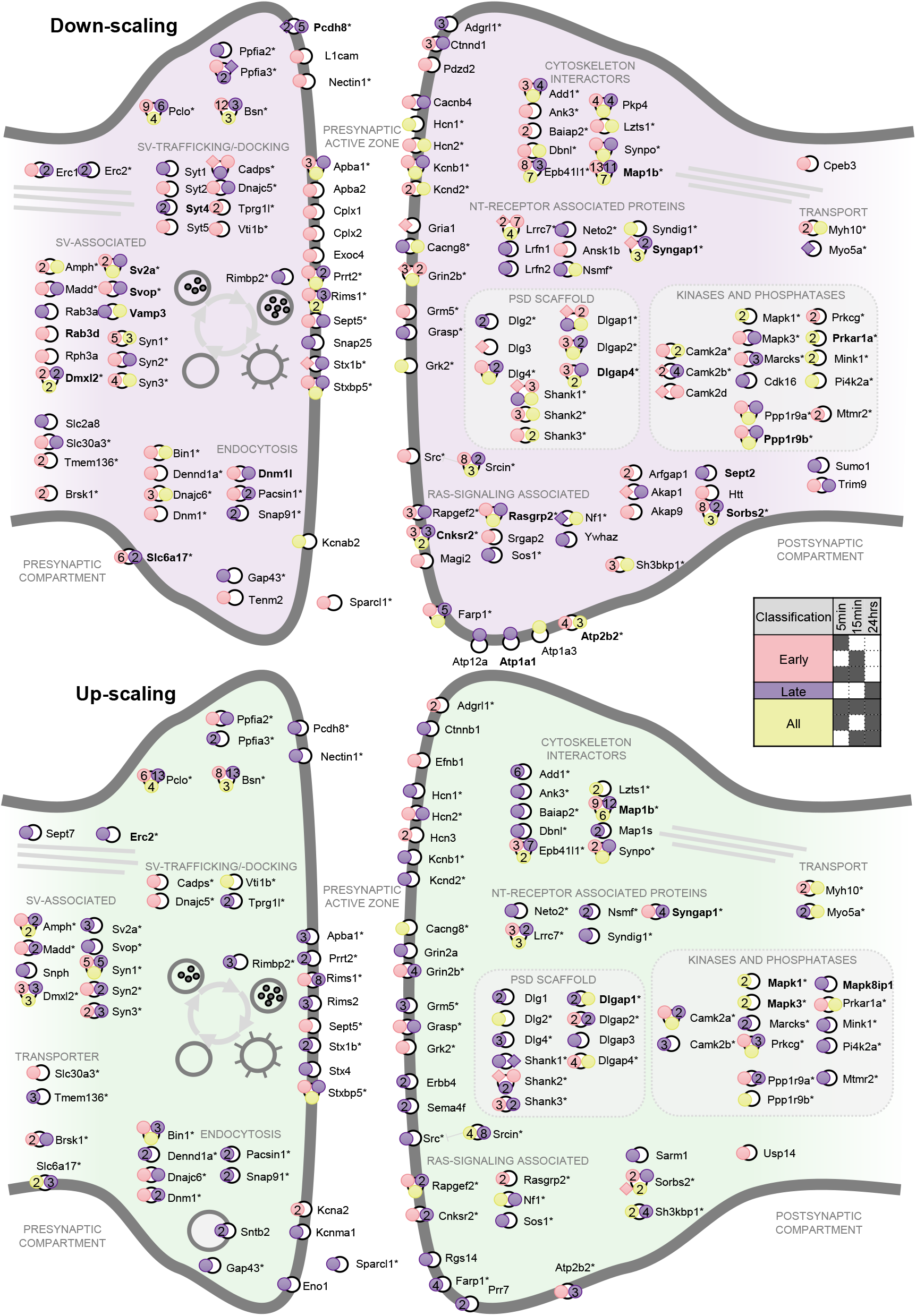
Activity-dependent protein phosphorylation in synaptic compartments. Differentially regulated phosphorylation sites on synaptic proteins during Bic-induced down-scaling (top) or TTX-induced up-scaling (bottom), displayed according to their synaptic localization. The regulated phosphorylation sites are displayed as circles on the respective protein and are colored according to their temporal categories (see legend). Proteins marked with an asterisk were phosphoregulated during both down- and up-scaling. Proteins highlighted in bold were regulated in protein abundance as well. In cases where multiple phosphopeptides covered the same phosphorylation site exhibited differences in their regulatory pattern, the classification of the singly phosphorylated peptide is displayed here.

In contrast to the highly-regulated groups/ phosphorylation hotspots, other phosphoproteins were exclusively regulated during one particular scaling phenotype. For example, phosphorylation events on the proteins complexin-1 (Cplx1) and complexin-2 (Cplx2) of the presynaptic active zone showed a significant change only during the early phase of Bic-induced down-scaling. In addition, Bic-induced phosphorylation of two postsynaptic proteins, synaptic adhesion-like molecule 1/2 (Lrfn2 and Lrfn1), known to interact with central proteins of the postsynaptic density (e.g. Dlg4, Gria1 and Grin1,Ko et al., 2006), was exclusively observed at 24 hrs. For TTX-induced up-scaling, three phosphorylation sites on Rims2, also a protein of the presynaptic active zone, were regulated only at 24 hrs.

Lastly, we compared the phosphoregulation we observed during our synaptic scaling experiments to phosphoregulated proteins detected in previously published phosphoproteomics experiments. In general, the phosphoregulated proteins in our dataset were much more extensive but overlapped with the phosphoproteins detected following other forms of synaptic plasticity (Figure S7A,B), including long-term potentiation (Li et al., 2016), depolarization (Kohansal-Nodehi et al., 2016) or mGluR-dependent LTD (van Gelder et al., 2020). Altogether, the phosphoproteins associated with synaptic scaling identified here captured between 60-75% of the phosphoproteins identified in these previous plasticity studies. In addition, we also identified ~40% of the phosphoproteins identified in a recent study examining the molecular timecourse of pathology associated with Alzheimer’s disease (Bai et al., 2020) (Figure S7B).

## Discussion

We generated a comprehensive, time-resolved map of the differential protein phosphorylation underlying homeostatic scaling in cultured cortical neurons. In response to the global activity manipulations that induce scaling, we detected both transient and persistent changes in over 3,300 phosphorylation events. We complemented our phosphoproteomic analyses with quantification of the total proteome and found a negligible contribution of protein abundance to the regulated phosphorylation events. Regulated phosphoproteins were significantly enriched for synaptic functions and cytoskeletal organization. As discussed below, a significant number of phosphoproteins were phosphorylated reciprocally during up- and downscaling, reflecting the opposing scaling polarity.

Previous high-throughput studies investigated protein phosphorylation as a regulatory element in sleep (Brüning et al., 2019; Diering et al., 2017), long-term potentiation (Li et al., 2016), depolarization (Engholm-Keller et al., 2019; Kohansal-Nodehi et al., 2016) or mGluR-dependent LTD (van Gelder et al., 2020). A recent study explored AD stage-associated investigations (Bai et al., 2020). While the overlap of phosphoregulation with different studies on phosphorylation in synaptic plasticity was extensive, phosphoregulation in homeostatic scaling encompassed a broader set of targets, indicated by the great number of phosphoproteins exclusively regulated during scaling. Our dataset is the first systematic study on protein phosphorylation during homeostatic synaptic scaling, examining phosphoproteomic changes that occur within minutes of activity-offset to those that are responsible for the implementation of scaling detected 24 hrs later, as discussed below.

A key feature of a homeostatic response system is an output set-point that is retargeted following an activity perturbation (Davis, 2006). To achieve homeostasis, an offset from the set-point needs to be sensed and coupled to effector mechanisms to bring about synaptic scaling. In principle, the early activity-sensitive phosphorylation events we detected could comprise offset “sensors” and the late phosphorylation events could comprise “effectors”.

We detected 282 and 1,039 regulated phosphorylation events unique to the early phase of up- or down-scaling, respectively; these modifications could be responsible for sensing the global change in activity. A subset of 110 events was shared between both up- and down-scaling. These common sensor events could detect a sign-dependent or sign-independent activity set-point offset. Indeed, we noted that 47% of these common early phosphoregulatory events were reciprocal in nature: manifest as either decrease or increase in phosphorylation, depending on the type of scaling. We noted that early time points, down-scaling was associated with many differential phosphorylation events than up-scaling. This could be due to rapid, Bic-evoked change in intracellular Ca^2+^ levels. Consistent with an early influx of Ca^2+^, we detected the largest increase in phosphorylation of the Camk2a Thr^286^ site directly after 5 min Bic-treatment. In addition, some Camk2a-associated phosphorylation events such as Ser^16^ of the cytoskeleton-interacting protein Stmn1 or the LTP-associated phosphorylation of Ser^831^ on the AMPA receptor subunit 1 (Gria1; exclusive phosphorylation; Lee, 2006) were uniquely detected during early downscaling. During early TTX-stimulation, the phosphorylation of Thr^286^ could not be reliably quantified as the phosphorylated peptide species was not abundant. We also detected Camk2a-associated sites among the shared, bi-directional sensor sites such as on the postsynaptic scaffold protein Shank3 (Ser^1511^; Dosemeci and Jaffe, 2010). Also the kinases ERK1/2, which can be secondarily activated by Ca^2+^-dependent mechanisms (Thomas and Huganir, 2004; Zhu et al., 2002), exhibited an early (5 min) Bic-evoked increase in both phosphorylation sites of its activation loop (Thr^203^/Tyr^205^; Thr^183^/Tyr^185^).

Phosphorylation events that were present during the late phase of synaptic up- or down-scaling represent potential effectors: modifications that are required to express the homeostatic response. We detected 912 and 681 potential effector events unique to the late phase of either up- or down-scaling, respectively. The number of common effector events shared between the opposite scaling groups was 149 with the majority (74%) exhibiting reciprocal, polarity-dependent regulation. Since synaptic scaling converges on changes of synaptic weight by changing the number of AMPA receptors in the postsynaptic membrane (O’Brien et al., 1998; Turrigiano et al., 1998), obvious effector proteins include glutamate receptors or receptor-associated molecules involved in receptor transport, surface retention or functional modulation. While we identified regulation of Gria1 in the early phase of activity-manipulation, we did not identify previously reported phosphorylation sites on the receptors during late phases of scaling, (e.g. Tyr^876^, after 48 h up-scaling; Yong et al., 2020). Nevertheless, our data indicate a pronounced phosphomodulation of glutamate-receptor associated and scaffolding proteins of the postsynaptic density. For example, we detected differential phosphorylation unique to the late phase of down-scaling on the proteins Lrfn1 or Lrfn2, known to induce clustering of excitatory proteins of the postsynaptic density (Ko et al., 2006) or involved in surface expression of AMPA receptor subunits (Seabold et al., 2008; Wang, 2006), respectively. Another example is the late up-scaling phosphoregulation of the protein Syndig1, which is known to interact with the AMPA receptor subunits Gria1 and Gria2 (Kalashnikova et al., 2010).

In contrast to time-sensitive phosphorylation events, we found that a considerable proportion (~25%) of initially regulated phospho-events were either continuously switched on or off (e.g. exhibited a persistent increase or decrease in phosphorylation). This set of phospho-events was associated with proteins of the postsynapse, involved in organization of the cytoskeleton and during down-scaling also in modulation synaptic transmission. The observed persistent phosphorylation differs from the proteomic regulation throughout homeostatic scaling, where little overlap in the identity of the newly-synthesized proteins of the early (2 hrs) and late phase (24 hrs) was observed (Schanzenbächer et al., 2018). Among the persistently regulated phospho-events, we found the Bic-evoked autophosphorylation sites of the kinases Camk2a (peak after 5 min Bic) and ERK1/2, suggesting both an early and late role for Ca^2+^-sensitive kinases. For example, we detected the persistent phosphoregulation of Ser^421^ of the transcription factor Mecp2 which has been reported as a critical factor during synaptic scaling (Blackman et al., 2012). Ser^421^ is known to be phosphorylated by Camk2a in an activity-dependent manner leading to induction of Bdnf transcription (Chen et al., 2003; Zhou et al., 2006). We found that Ser^421^ of Mecp2 was indeed phosphorylated during all time points following Bic-treatment and we detected a concomitant upregulation of Bdnf after 24 hrs Bic stimulation. Interestingly, a role for persistent kinase activity has been previously reported in several *in vivo* studies suggesting a role in behavioral plasticity or long-term memory. For example, continuous hippocampal Camk2a autophosphorylation was observed up to 20 hrs after inhibitory avoidance training (Bambah-Mukku et al., 2014). In modeling studies, persistent autophosphorylation of Camk2a has been proposed to contribute to long-term information storage (Lisman and Goldring, 1988; Lisman and Raghavachari, 2015). Moreover, autophosphorylation of ERK1/2 kinases in long-term memory had been observed during spatial learning experiments with rodents (Blum et al., 1999; Selcher et al., 1999) and persistence in ERK1/2-activity was proposed in modeling studies of memory maintenance (Smolen et al., 2008).

Mechanistically speaking, the question arises if the phenotypically opposite poles of up- and down-scaling are mediated by reciprocal regulation of common proteins? Proteomic analyses have identified commonly regulated proteins in a bi-directional manner during up- and down-scaling, but mostly divergence, suggesting the potential involvement of unique cellular pathways (Dörrbaum et al., 2020; Schanzenbächer et al., 2016). In this study, we identified 219 phosphorylation events that strictly follow a bi-directional pattern reflecting the opposing scaling polarities. We found that persistent phosphorylation events associated one scaling paradigm were persistently and reciprocally regulated in the opposing scaling paradigm. Indeed, the majority (Bic: 68%; TTX: 85%) of the persistent phosphorylation events were associated with bidirectional regulation when directly compared to one another in cluster analysis. The reciprocal and persistent phosphorylation events that we observed are predominantly associated with activity modulation of Camk2a or MAP kinases ERK1/2: autophosphorylation sites of these kinases and also the differential kinase activity inferred from KSEA largely matched the same bi-directional, continuous pattern which was also verified by phospho-specific antibodies. The results of these analyses suggest persistence in catalytic kinase activity, however, a slower rate of dephosphorylation by phosphatases could also contribute. Reciprocal phosphorylation might also represent a polarity-specific tagging component allowing capture and/or retention of plasticity-related mRNA or proteins driving downstream mechanism leading to scaling of corresponding polarity. Examples of phosphorylation sites that could serve as bi-directional switches are Ser^422^ or Ser^516^ on Dlg4 or Arhgef7, postsynaptic proteins involved in receptor-trafficking and signaling, respectively. In the late phase of scaling, phosphorylation of the Ser^418^ of Dlg4 in the same reciprocal nature as Ser^422^ was also observed. With respect to protein structure, both sites are directly localized between the PDZ-3 and SH3 domain of Dlg4. These domains known to enable the interaction with a broad spectrum of proteins (Kim and Sheng, 2004) and affect molecular association/ protein localization (Pedersen et al., 2017). Overall, many synaptic proteins appeared as phosphorylation hotspots where phosphoregulation of different temporal categories converged, demonstrating that one protein can have multiple roles throughout the expression of the homeostatic response.

Taken together, our findings highlight protein phosphorylation as a major molecular driver following global activity-manipulations over time scales ranging from minutes to a day. Phosphomodulation during scaling induction was not only achieved by time-limited phosphorylation, but persistent and strictly bi-directional phosphorylation was also shown to play a prominent role in both phenotypes of homeostatic plasticity connecting them mechanistically. In support of our data, we detected differential regulation of proteins as well as phosphorylation events that had already been reported in context of synaptic plasticity, but also identified new phosphoregulated candidates. The broad detection of distinct phosphorylation profiles provides insights into the fundamental processes that underlie activity sensing and scaling manifestation. These data thus provide resource for further candidate-based investigations into the role of long-lasting and/or bi-directional phosphorylation during homeostatic scaling and other forms of plasticity.

## Acknowledgements

We thank I. Bartnik, N. Fuerst, A. Staab and C. Thum for the preparation of primary cell cultures, A. Dörrbaum for valuable advice on the statistical analyses and F. Rupprecht for MS maintenance and assistance with data acquisition. E.M.S. is funded by the Max Planck Society, an Advanced Investigator award from the European Research Council (grant No. 743216), DFG CRC 1080: Molecular and Cellular Mechanisms of Neural Homeostasis, and DFG CRC 902: Molecular Principles of RNA-based Regulation.

## Author Contributions

KD designed, conducted and analysed experiments and wrote the paper. JDL and EMS designed experiments, supervised the project and co-wrote the paper.

## Methods

### Preparation and culture of primary cortical cells

Dissociated cortical neurons were prepared and maintained as previously described for hippocampus neurons (Aakalu et al., 2001). Cortices from postnatal day one old rat pups (RRID:RGD_734476; strain Sprague-Dawley) were dissected, dissociated by incubating with L-cysteine-papain solution at 37°C and plated onto 10 cm Petri dishes (MatTek, Ashland, MA) previously coated with poly-D-lysine. Cultured cells were kept in Neurobasal-A medium (Invitrogen, Carlsbad, CA) supplemented with B-27 (Invitrogen) and Glutamax (Invitrogen) at 37°C and 5% CO2 for 19-20 days.

### MS-sample preparation and phosphopeptide enrichment

Nine dishes (3 million cells/ dish) were prepared for each experiment. Three dishes each were treated with either 20 μM Bicuculline, 1 μM Tetrodotoxin or no drug (control) for 5 min, 15 min or 24 hrs. Afterwards, the cells were harvested by briefly washing with ice-cold DPBS (Invitrogen) supplemented with protease inhibitor cocktail (cOmplete EDTA-free; Roche, Basel, Switzerland) and phosphatase inhibitors (PhosStop; Roche), followed by scraping and pelleting by centrifugation.

Cell pellets were lysed using lysis buffer (5% SDS, 25 mM Tris, pH 7.55, supplemented with protease and phosphatase inhibitor), and then disrupted with a pipette and four sonication cycles for 30 s. Lysates were incubated with Benzonase (1 μl; 250 units/mL stock solution; Sigma, St. Louis, MO) for 10 min at room temperature. To clear debris from the samples, they were centrifuged for 8 min at 13,000 x g. Protein concentration was determined by a BCA assay (ThermoFisher Scientific, Waltham, MA). The samples were diluted 1:5 prior to the assay to minimize interference of the high detergent concentration.

For bottom-up MS analysis, protein digestion was performed according to an adapted version of the suspension trapping protocol as described by the manufacturer (S-Trap, ProtiFi, Huntington, NY). In brief, 350 μg of protein in lysis buffer was reduced by DTT addition in a final concentration of 20 mM for 10 min. Then proteins were alkylated using iodoacetamide at a final concentration of 40 mM and incubated for 30 min at room temperature in the dark. Afterwards, the sample was acidified by addition of phosphoric acid to a final concentration of 1.2%. Binding buffer (90% methanol, 50 mM TRIS, pH 7.55) was added in a 1:7 lysate to buffer ratio. The mixture was loaded onto the S-Trap filter (mini) by centrifugation for 30 s at 4,000 x g in 450 μl-steps and washed with 400 μl binding buffer for four times. Sequencing-grade trypsin (Promega, Madison, WI) was added in 150 μl digestion buffer (40 mM ammonium bicarbonate) in an enzyme-to-protein ratio of 1:50. The protease buffer was briefly (1-3 s) spun into the trap; solution passing the filter was re-added on top. Digestion was carried out overnight at room temperature under gentle agitation and in a humidified chamber to prevent filters from drying out. To elute peptides, the filter was rinsed in three consecutive steps by centrifugation at 1,000 x g for 60 s starting with 80 μl digestion buffer and two 80 μl washes with 0.2% formic acid (FA) in MS grade water.

After digestion, the peptides were desalted using C_18_-SepPak columns (50 mg sorbent; Waters, Milford, MA) as previously described (Schanzenbächer et al., 2016). Desalted peptides were separated for analysis of the total proteome (5% v/v) and subsequent enrichment for phosphorylated peptides (95% v/v). All samples were dried *in vacuo* using a Speed Vac (Eppendorf, Hamburg, Germany) at room temperature and stored at −20°C until LC-MS analysis or further use. Enrichment for phosphorylated peptides was performed using Titanium dioxide beads (TiO_2_; kit: #432993; ThermoFisher Scientific) as described in the manufacturer’s protocol. Eluted peptides were dried *in vacuo* at room temperature and stored at −20°C until LC-MS analysis.

Each experiment was carried out in four independent biological replicates.

### LC-MS/MS analysis

Dried peptides or phosphorylated peptides were reconstituted in 5% acetonitrile (ACN) with 0.1% FA or 2% ACN with 0.1% FA, respectively. Peptides were loaded onto a C18-PepMap 100 trapping column (particle size 3 μm, L = 20 mm) and separated on a C18-EasySpray analytical column (particle size = 2 μm, ID = 75 μm, L = 50 cm, ThermoFisher Scientific) using a nano-HPLC (Dionex U3000 RSLCnano). Temperature of the column oven was maintained at 55°C.

Trapping was carried out for 6 min with a flow rate of 6 μl/min using loading buffer (100% H2O with 0.05% triflouroacetic acid). Peptides were separated by a gradient of water (buffer A: 100% H2O and 0.1% FA) and acetonitrile (buffer B: 80% ACN, 20% H2O and 0.1% FA) with a constant flow rate of 300 nL/min. The gradient for unmodified peptides went from 4% to 48% buffer B in 180 min, the gradient for phosphorylated peptides from 4% to 30% buffer B in 110 min and to 45% buffer B in 10 min. All solvents were LC-MS grade and purchased from Riedel-de Häen/Honeywell (Seelze, Germany).

Eluting peptides were analyzed in a data-dependent acquisition mode on a Fusion Lumos mass spectrometer (ThermoFisher Scientific) coupled to the nano-HPLC (Dionex U3000 RSLCnano) by an EASY Spray ESI source. MS1 survey scans were acquired over a scan-range of 350 to 1400 mass-to-charge ratio (m/z) in the Orbitrap detector (resolution (R) = 120k, automatic gain control (AGC) = 2e5 and maximum injection time: 50 ms,). Sequence information was acquired by a “top speed” MS2 method with a fixed cycle time of 2 s for the survey and after MS/MS scans. MS2 scans were generated from the most abundant precursors with a minimum intensity of 5e3 and charge states from two to five. Selected precursors were isolated in the quadrupole using a 1.4 Da window and fragmented using higher-energy C-trap dissociation (HCD) at 30% normalized collision energy. For MS2, an AGC of 1e4 and a maximum injection time of 300 ms were used. Resulting fragments were detected in the ion trap using the rapid scan rate (AGC = 1e4, maximum injection time = 300 ms). Dynamic exclusion was set to 30 s with a mass tolerance of 10 parts per million (ppm). All LC- and MS-parameters are listed in the supplementary Table S1. Each sample was measured in triplicate LC-MS/MS runs.

### MS-data processing

MS raw data were processed using the MaxQuant software (ver. 1.6.6.0; RRID:SCR_014485; Cox and Mann, 2008) with customized parameters for the Andromeda search engine. For all searches, spectra were matched to the *Rattus norvegicus* database downloaded from the UniProtKB (RRID:SCR_004426; Proteome_ID: UP000002494; downloaded on 23 August 2019), a contaminant and decoy database. Tryptic peptides with a minimum length of seven amino acids and a maximum of two missed cleavage sites were included. Precursor mass tolerance was set to 4.5 ppm and fragment ion tolerance to 0.5 Da. Carboxyamidomethylation of cysteine residues was set as a static modification. Acetylation (Protein-N-term.) and oxidation of methionine residues were assigned as variable modifications. Analysis of the phosphoproteome included the assignment of phosphorylation of serine, threonine and tyrosine residues as variable modification. With the use of a decoy strategy, a false discovery rate (FDR) below 1% at protein, peptide and modification level was applied. The “match between runs” option was enabled (matching time window = 0.7 min, alignment time window = 20 min). Only proteins identified by at least one unique peptide were considered for further analysis. Label-free quantification of proteins was performed by pair-wise ratio determination using at least two common peptides in at least three consecutive full scans (Cox et al., 2014). For all details on the parameters for raw-data processing in MaxQuant see the supplementary Table S2.

All data associated with this manuscript have been uploaded to the PRIDE repository and are available with the dataset identifier PXD021834 (RRID:SCR_003411; Vizcaíno et al., 2013).

### Proteomics data postprocessing and statistical analysis

Four independent biological replicates measured in triplicates were processed using the Perseus software (ver. 1.6.2.3; RRID:SCR_015753; Tyanova et al., 2016). MaxQuant protein results (total proteome: proteinGroups.txt) and results of the phosphorylation site table (phosphoproteome: phospho(STY)sites.txt) were filtered for contaminants and reverse (decoy) database hits. The phosphoproteomic data were further filtered for localization probability of the phosphorylated residue greater than 75% to trace the modification to one particular residue. Then the phosphorylation site table was rearranged according to multiplicity, i.e. an expansion of entries so that the number of phosphorylation sites per peptide was formatted as separate rows. Species of this modified table were called phosphorylation events, since there can be more than one entry for a particular phosphorylation site, e.g derived from a singly, doubly or triply phosphorylated peptide species. Protein LFQ intensities and phosphorylation event intensities were log_2_-transformed and normalized according to the sample’s median intensity.

For quantitative comparisons at each time point (5 min, 15 min and 24 hrs), only treatment-control pairs of proteins or phosphorylation events, which could be quantified in all four biological replicates in at least one technical replicate, were considered for further statistical analysis. Additional valid value filtering required 50% valid entries across each pair-wise data matrix. We used the lme4 package in the statistical computing software R to perform a linear mixed effect analysis (RRID:SCR_015654; Bates et al., 2015). Differential regulation of treatment versus control was investigated using a model where the treatment was considered the fixed effect in question and the biological replicate was entered as a possible random effect, similar to previously reported strategies (Dörrbaum et al., 2020). P-values were calculated by likelihood ratio tests of the model including the effect of interest against the model without it. To correct for multiple testing, Benjamini-Hochberg correction was applied with an FDR cut-off < 0.01 (Benjamini and Hochberg, 1995).

### Bioinformatic tools for phosphoproteomics analyses

#### Hierarchical clustering

For hierarchical clustering analysis, log_2_-tranformed fold changes of treatment against control intensities of significantly regulated phosphorylation events were used. The particular segmentation of the data in the clusters was stated in the corresponding figure legends. Distance measure to cluster the rows was Euclidean distance. Clustering was performed using the “ward.D2” agglomeration algorithm. Visualization using this metrics was done using the pheatmap package in the statistical computing software R (RRID:SCR_016418; Kolde, 2019).

#### Kinase substrate enrichment analysis

To match differentially regulated protein phosphorylation sites to their reported kinases, kinase substrate enrichment analysis was carried out using the KSEAapp package in the statistical computing software R (Wiredja et al., 2017). Site-specific information on the kinase-substrate pairs was downloaded from PhosphositePlus.org database (on 5^th^ November 2019; RRID:SCR_001837; Hornbeck et al., 2015) and filtered for species-specific entries to *Rattus norvegicus*. NetworKIN predictions were excluded. For survey, a minimum number of one substrate of a kinase was set. Kinase z-scores representing the normalized score of each kinase weighted by the number of identified substrates were calculated. Multiple hypothesis testing to asses a p-value for the kinase z-score was corrected using the Benjamini-Hochberg method (Benjamini and Hochberg, 1995). The log_2_-transformed fold change of this site is reported and if a particular substrate phosphorylation site was detected across multiple phosphorylated peptide, its average is used in the algorithm.

#### Phosphoprotein-Interaction Map

Proteins harboring a phosphorylation event that was regulated in four out of six conditions (Figure 3) were analyzed for protein interactions using their gene identifiers and the STRING database (RRID:SCR_005223; Franceschini et al., 2013). Interactions with a confidence > 0.4 (combined score) were included in the analysis. Interactions solely based on textmining as a source were excluded. Networks were exported and visualization was performed using the software Cytoscape (ver. 3.7.2; RRID:SCR_003032; Shannon, 2003).

### Comparison of proteome remodeling during synaptic scaling

The differentially regulated proteome characterized in this study was compared to results of another system-wide proteomics investigation of homeostatic scaling. For this purpose, we analyzed the data published by Doerrbaum *et al*. where primary-cultured hippocampal neurons were subjected to Bic or TTX treatment for one, three and seven days or left untreated (Dörrbaum et al., 2020). For direct comparison, only data of the control condition and the 24 hr treatments were considered. Quantification of proteins needed to have at least one valid treatment-control pair in all three biological replicates. For statistical evaluation, a linear mixed effects model was used as described in Dörrbaum et al., 2020. In brief, treatment was set as fixed effect in question, biological replicate and peptide identity nested into biological replicate were set as random effects. This resulted in 150 and 239 proteins that significantly changed in abundance in response to Bic or TTX treatment (Benjamini-Hochberg correction, FDR < 0.01, Benjamini and Hochberg, 1995). Proteins quantified in both studies exhibiting significant regulation at least one or both of them were used for correlation analysis.

### Synaptic Phosphoproteome

Proteins of the synaptic phosphoproteome (Figure 5) were selected if they matched a synaptic GO term for the cellular compartment (GOCC) and carried at least one significantly regulated phosphorylation site. Localization within the synapse was derived from the UniProtKB database information on subcellular localization. If there was more than one phosphorylation event for a particular phosphorylation site on a protein, the site-specific, temporal categorization (early, late or all) was derived from the species with the lowest multiplicity to achieve highest residue resolution. Phosphorylation events which were termed and highlighted as “exclusive” for a certain time point were excluded from statistical analysis as they were quantified reliably only in one treatment group but not in the control or other treatment groups. Nonetheless, the peptides were visualized in the overview (depicted as a square) as this subset were reliably quantified in all four biological replicates of the condition in question and not detected in all biological replicates of any other possible conditions.

### Comparison to other phosphoproteomics studies

To compare phosphoregulation associated with synaptic plasticity (van Gelder et al., 2020; Kohansal-Nodehi et al., 2016; Li et al., 2016), we extracted the significantly phosphoregulated species of the studies reported in different mass-spec based, high-throughput analyses. Significance was determined by the statistics and parameters reported in each study. We finally compared phosphomodulation on the level of phosphoregulated proteins by matching the studies via a species-neutral ID, gene names (Figure S7A,B).

### Western blot analysis

Primary cortical cultures were prepared and maintained as described above. After, 19 DIV, cells were incubated with either 20 μM Bicuculline, 1 μM Tetrodotoxin or no drug (control) for 5 minutes or 24 hrs. Cell lysates were prepared as described and equal protein amounts were loaded onto 4% to 12% Bis-Tris NuPAGE gels (ThermoFisher Scientific). After electrophoreses, proteins were transferred to a PVDF membrane (Immobilon-FL; Merck Millipore, Billerica, MA). Immunoblotting was performed with primary antibodies against Camk2a phosphorylated at pT^286^ (1:1000, Cell Signaling Technology Ref.: 12716), total Camk2a (1:1000, Invitrogen Ref.: 13-7300), Mapk1/ Mapk3 phosphorylated at pT^183^-pT^185^/ pT^203^-pY^205^ (1:1000, Cell Signaling Technology, Ref.: 4370) or total Mapk1/ Mapk3 (1:1000, Cell Signaling Technology, Ref.: 4696). Secondary antibodies, anti-mouse (1:15,000, Ref.: 926-68020) and anti-rabbit (1:15,000, Ref.: 926-3211) were purchased from LI-COR (Lincoln, NE). Densiometric quantification was conducted using LI-COR Image Studio Lite (RRID:SCR_013715). All experiments were performed in three independent biological replicates.

## Supplemental Figures

**Figure S1 –.**
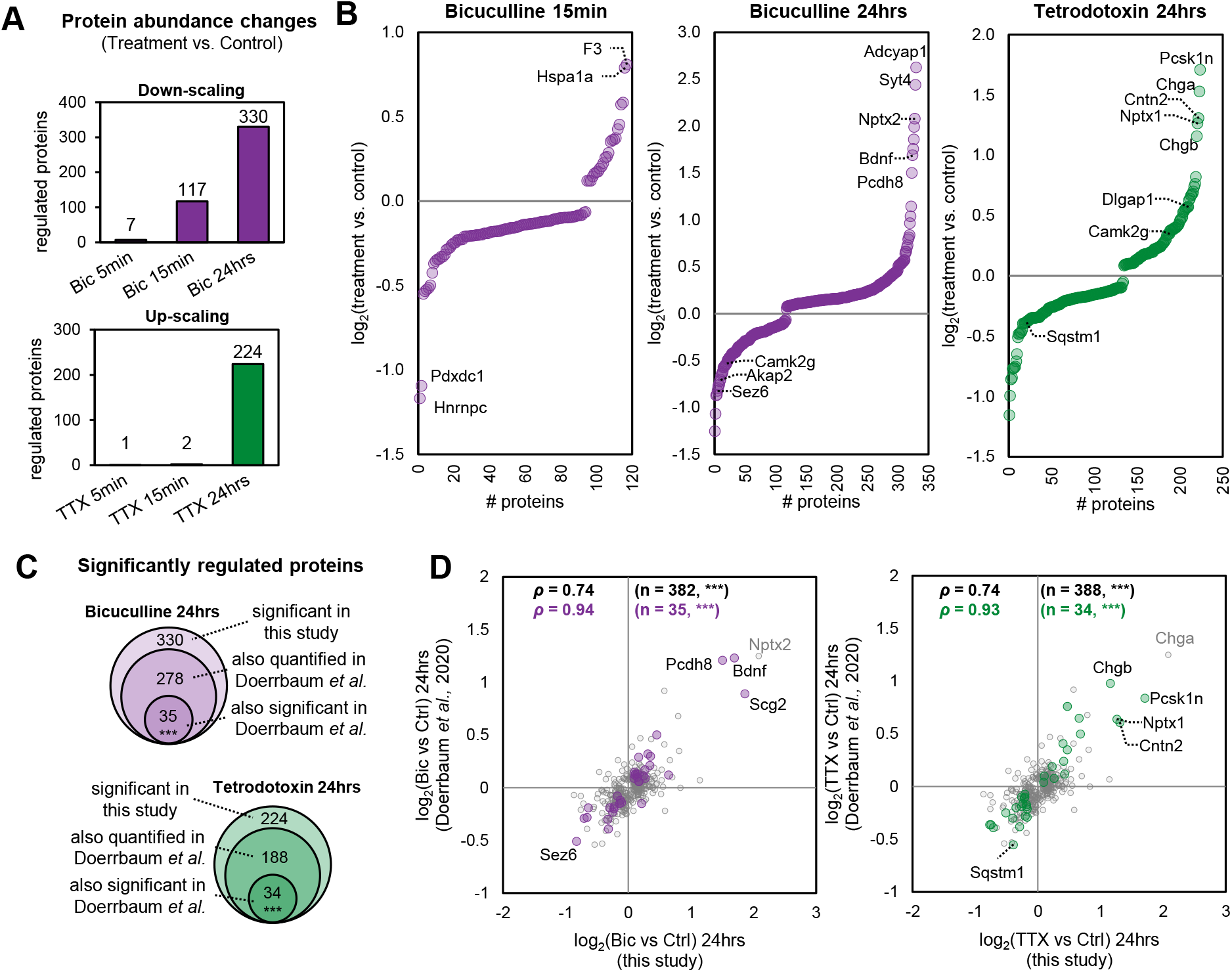
Remodeling of the cortical proteome during homeostatic synaptic scaling. (A) Bar chart depicts the number of significantly regulated proteins at each time point of Bic or TTX treatment compared to the control group (Benjamini-Hochberg correction; FDR<0.01). (B) Rank-ordered log_2_ fold changes of the regulated proteins (treatment vs. control) for the three time points showing the greatest changes in protein abundance: Bic treatment for 15 min and 24 hrs and TTX treatment for 24 hrs. (C) Comparison of the significantly regulated proteins from this experiment to Doerrbaum *et al*. (see Methods), 278 of the 330 proteins regulated by Bic and 188 of 224 regulated by TTX were also identified in their dataset. Significantly more proteins were regulated in experiments of both studies after 24 hours of Bic and TTX treatment than by chance (Fisher’s exact test; *** p <0.001; fold enrichment (FE): FE_Bic_ = 3.4, FE_TTX_ = 2.9). (D) Pearson correlations of the log_2_ fold changes (treatment vs. control) of the proteins quantified in both studies were significant (*** p <0.001): The Pearson correlation coefficient (ρ) of the overlapping proteins significantly regulated after 24 hours Bic or TTX treatment in at least one of the studies was 0.74 in both cases (n_Bic_ = 382, n_TTX_ = 388; grey). The Pearson correlation coefficient of the overlapping proteins exhibiting significant regulation in both studies was 0.94 or 0.93, respectively (n_Bic_ = 35, n_TTX_ = 34; purple/ green)

**Figure S2 –.**
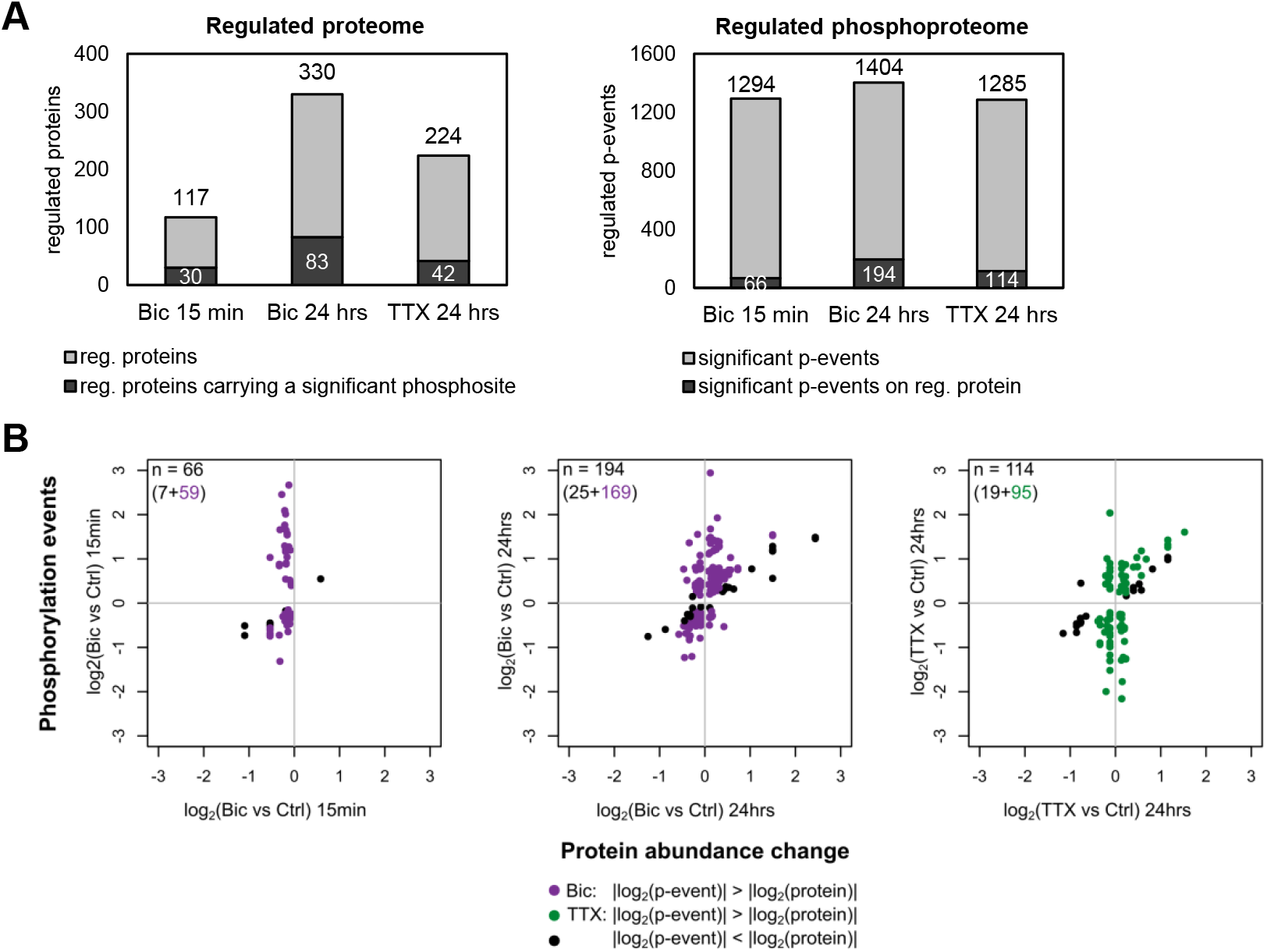
Regulation of proteome remodeling and phosphorylation. (A) Bar plot (left) shows the significantly regulated proteins after Bic treatment for 15 min and 24 hrs or TTX treatment for 24 hrs, highlighting the proportion of regulated proteins that carry at least one significantly regulated phosphorylation site in the same condition in a dark shade. Bar plot (right) displays the significantly regulated phosphorylation events after Bic treatment for 15 min and 24 hrs and TTX treatment for 24 hrs, highlighting the proportion of events that are located on a protein exhibiting significant abundance changes in the same condition. (B) Scatterplot shows the log_2_ fold changes of the overlapping subset of regulated phosphorylation events versus the log_2_ fold changes of the corresponding regulated protein. In cases where the change of the phosphorylation event was greater than that of the protein it is located on, the dots are highlighted by color. Black dots indicate a greater change in protein abundance. The majority of phosphorylation events were found to be much more dynamic in all three conditions. The amplitude of regulation on phosphorylation level exceeded those on proteome level. This indicates the activity-dependent changes in phosphopeptide abundance observed in our dataset mostly arise from differential phosphorylation rather than proteome remodeling.

**Figure S3 –.**
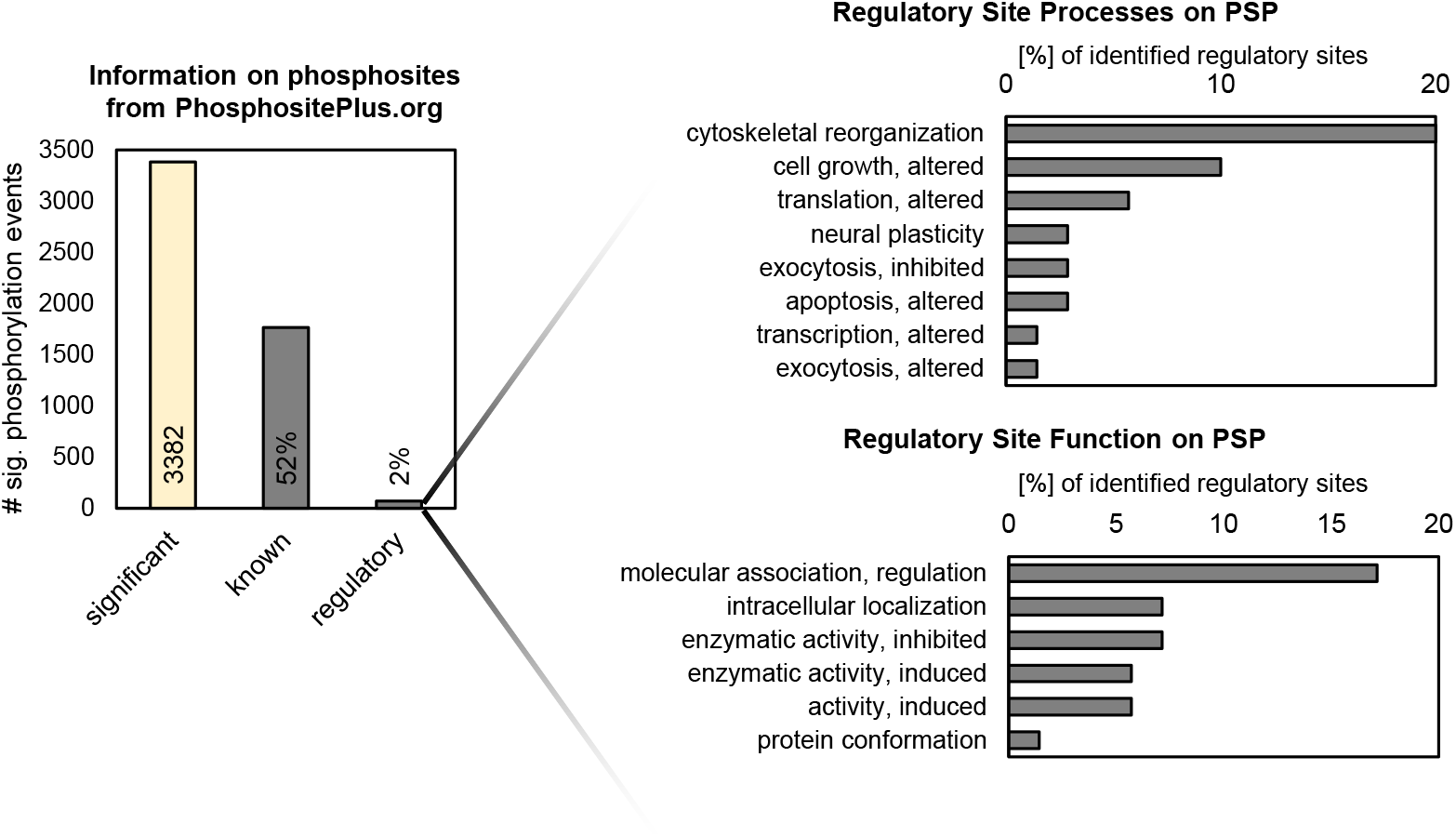
Annotation of regulated phosphorylation events across all experimental groups. Significant change in phosphorylation was detected for 3,382 different phosphorylation events regulated along all experimental conditions (Benjamini-Hochberg correction; FDR <0.01). Information on the sites was assigned using the annotations of the PhosphoSitePlus database (Hornbeck et al., 2015). About 50% of the activity-dependent events had been identified in previous experiments (known) and 2% had a regulatory annotation (regulatory). On the left, the terms assigned to characterize the process (top) or function (bottom) of the regulatory sites are listed.

**Figure S4 –.**
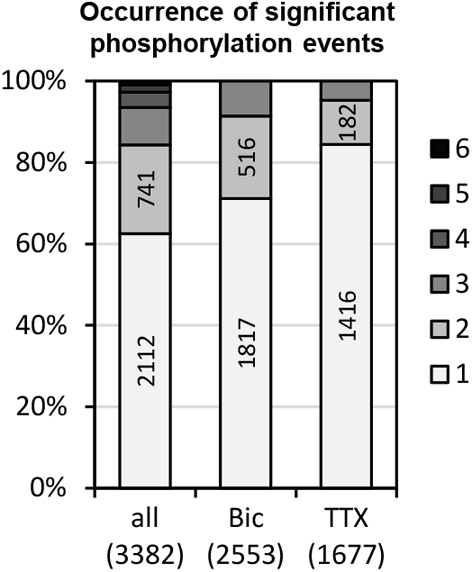
Significant phosphoregulation across time points. Bar chart showing an overview of the occurrence of significantly regulated phosphorylation events across all time points. The first bar summarizes the occurrence of significant regulation of a particular site with respect to all experimental conditions (Bic or TTX: 5 min, 15 min, 24 hrs). 2,112 (62%) of all regulated phosphorylation events were associated with a single time point. The second and third bar summarize the occurrence of significant regulation during up- or down-scaling (n/3). The majority of significant phosphorylation events are regulated in only one of the conditions (down-scaling: 72% and up-scaling 84%)

**Figure S5 –.**
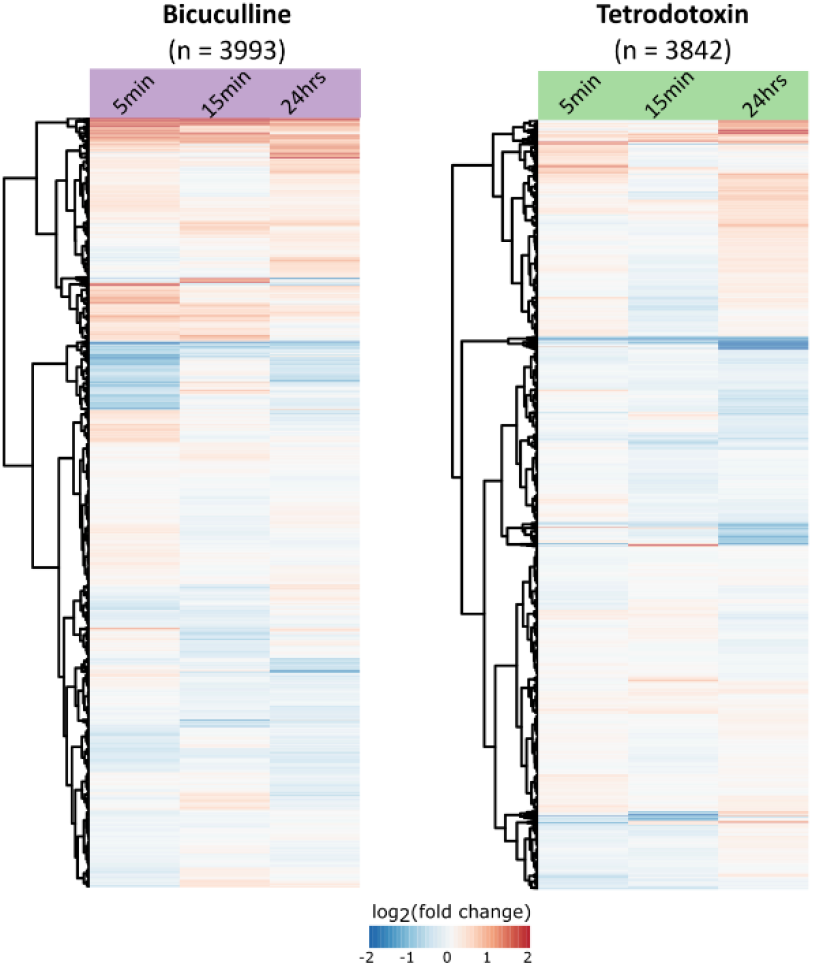
Global overview on the activity-dependent phosphoproteome. Temporal profile of the total data landscape of Bic-evoked (left) or TTX-evoked (right) phosphorylation events. Hierarchical clustering (Euclidean distance) was performed on the log_2_ fold change of phosphorylation event intensities (treatment vs. control) quantified at each time point of Bic and TTX treatment without applying any filter for significant regulation. The clustering yielded a heatmap with rather diffuse regulation (red: increase in phosphorylation, blue: decrease in phosphorylation), but trending towards persistent regulation which clearly emerged by selection of significant regulators (Figure 2D).

**Figure S6 –.**
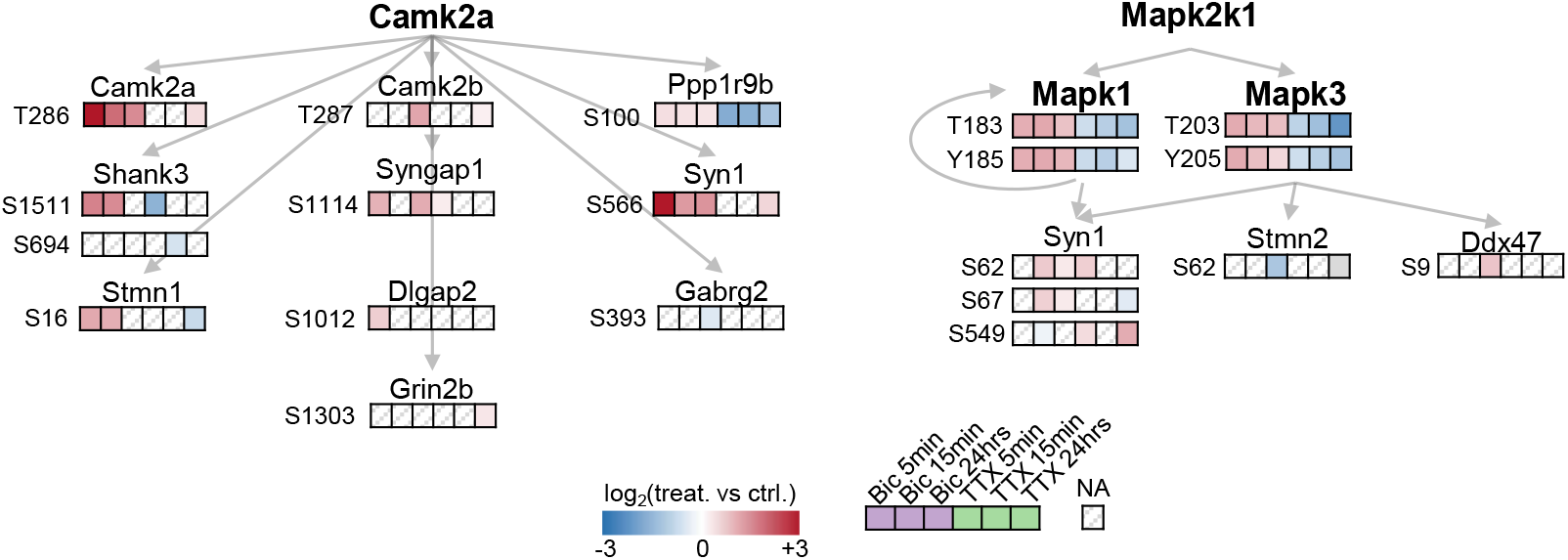
Phosphorylation status of Camk2a and Mapk1/3 associated phosphorylation sites. Phosphorylation status of significantly regulated Camk2a and Mapk1/3 target sites as reported in the rat-specific kinase-substrate database (PhosphoSitePlus) and identified in our dataset. The log_2_ fold change of the phosphoregulation (treatment vs. control) is highlighted by color. In case of multiple peptides covering the same phosphorylation site, the log_2_ fold change of a particular site was calculated as mean of the events. If the regulation was not significant or if there was no treatment-control pair identified, the box is marked as missing value (NA).

**Figure S7 –.**
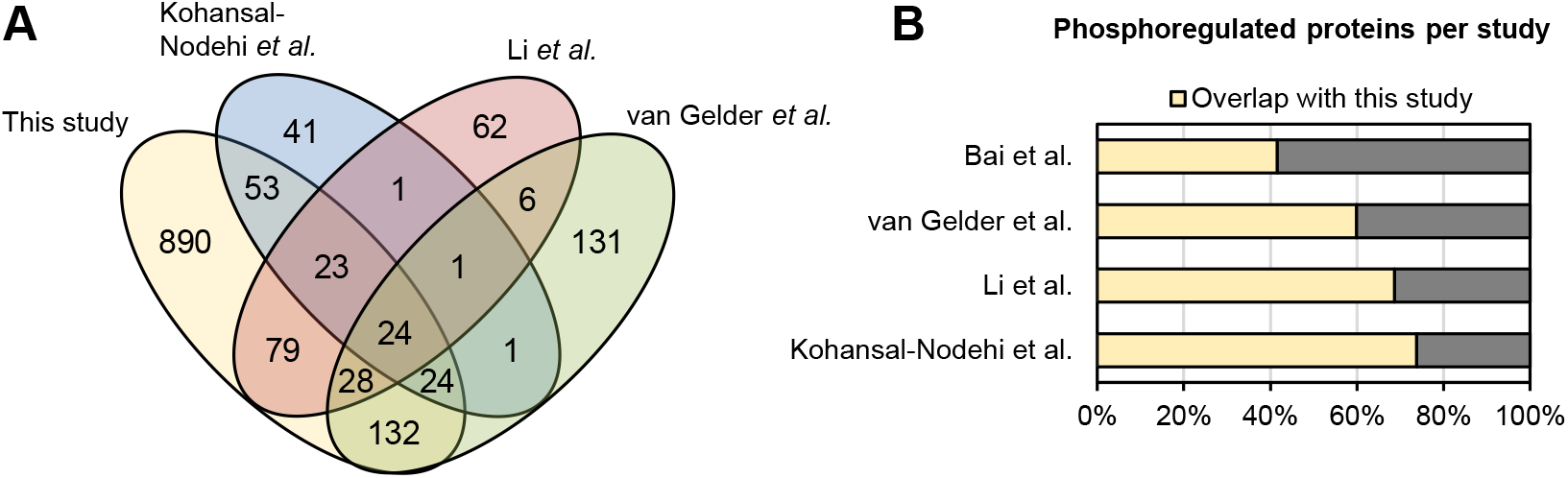
Comparison of phosphoregulated proteins with other phosphoproteomic studies of synaptic plasticity or neuronal function. (A) Venn diagram highlighting the overlap of proteins that carried at least one significantly regulated phospho-site originating from three different studies focusing on different aspects of synaptic plasticity, such as LTP (Li et al., 2016), depolarization (Kohansal-Nodehi et al., 2016) or mGlur-dependent LTD (van Gelder et al., 2020) with our dataset (for details see Methods). (B) Barplot indicates the significantly phosphoregulated proteins of each study presented in (A) and including a human AD stage-associated investigation (Bai et al., 2020). Each bar indicates the entity of regulated phosphoroteins per study, the colored fraction highlights the shared phosphoregulated proteins with our dataset (see Methods).

## Supplemental Tables

Supplementary table S1: Full list of LC- and MS-settings for the proteomic and phosphoproteomic analyses.

Supplementary table S2: Full list of parameters of MaxQuant search engine for MS-data processing of proteomic and phosphoproteomic analyses.

Supplementary table S3: Significantly regulated phosphorylation events during homeostatic scaling (related to Figure 1C,D).

Supplementary table S4: Significantly regulated proteins during homeostatic scaling (related to Figure S1).

Supplementary table S5: Classification of time-sensitive phosphorylation events during the different phases of homeostatic scaling (related to Figure 2A).

Supplementary table S6: GO term overrepresentation for subset of persistent phosphoproteins during up- and down-scaling (related to Figure 2E).

Supplementary table S7: Kinase substrate enrichment analysis for the separate time points of up- and down-scaling (related to Figure 3 and S6).

Supplementary table S8: List of synaptic phosphorylation events (related to Figure 5).

